# Enhancing Tissue-Specific Antiviral Immunity to Disrupt Arbovirus Transmission by Mosquitoes

**DOI:** 10.1101/2025.04.04.647234

**Authors:** Shengzhang Dong, Jessica Ciomperlik-Patton, Yuxuan Zhao, Yuemei Dong, Kevin M. Myles, George Dimopoulos

**Affiliations:** W. Harry Feinstone Department of Molecular Microbiology and Immunology, Bloomberg School of Public Health, Johns Hopkins University, 615 N. Wolfe Street, Baltimore, MD 21205-2179, USA; Department of Entomology and AgriLife Research, Texas A&M University, College Station, TX 77843-2475, USA

**Keywords:** *Aedes aegypti*, siRNA pathway, *Dicer2*, arboviruses, transgenic mosquitoes, salivary gland, fat body, midgut, antiviral mechanism

## Abstract

Arboviruses, including dengue virus (DENV), Zika virus (ZIKV), and chikungunya virus (CHIKV), pose a significant global health and economic burden, with *Aedes aegypti* serving as their primary vector. Arbovirus infection in *Ae. aegypti* progresses sequentially through the midgut (MG), carcass (CA), and salivary glands (SG), with each tissue exhibiting distinct antiviral responses. Here, we investigate tissue-specific antiviral mechanisms, focusing on the small interfering RNA (siRNA) pathway in SGs. Our results reveal that SGs possess weaker antiviral defense and are more susceptible to arboviral infection compared to MGs and CAs. Notably, overexpression of *Dicer2* (*Dcr2*), a key component of the siRNA pathway, in SGs leads to a significant decrease in arboviral replication. Conversely, *Dcr2* overexpression in fat bodies, the primary tissue in CAs, only moderately suppresses DENV2 infection and has no notable effect on Mayaro virus (MAYV) infection. Remarkably, the simultaneous overexpression of *Dcr2* in both MGs and SGs enhances antiviral activity, effectively blocking the transmission of multiple arboviruses. These findings reveal the tissue-specific dynamics of mosquito antiviral immunity and underscore the potential for targeting SG-specific immunity to disrupt arbovirus transmission, providing a promising approach for controlling mosquito-borne diseases.

## Introduction

*Aedes aegypti* is a key vector for several human viral pathogens, including dengue virus (DENV), Zika virus (ZIKV), and chikungunya virus (CHIKV). These arthropod-borne viruses (arboviruses) circulate between human hosts and mosquito vectors, contributing to significant global economic and health challenges (Bhatt et al., 2013; Weaver et al., 2018). Existing strategies to control mosquito-borne diseases have proven inadequate due to factors like insecticide resistance, climate change, and the limited availability of effective vaccines for many arboviruses (Achee et al., 2015; Ebi and Nealon, 2016; Moyes et al., 2017; Vontas et al., 2012). As a result, complementary approaches, such as genetic control methods, have been proposed to combat the increasing prevalence of mosquito-borne diseases (Alphey et al., 2013; Caragata et al., 2020; Kefi et al., 2024; Wang et al., 2021).

Upon infection, arboviruses initially infect the mosquito midgut, where they replicate in epithelial cells. The virus then disseminates into the mosquito’s hemocoel, infecting tissues such as fat bodies and hemocytes, before reaching the salivary glands (SGs), the last mosquito tissue to be infected by an arbovirus. Once established in SGs, viruses are released into mosquito’s saliva and transmitted to a human host during blood-feeding (Franz et al., 2015; Lee et al., 2019; Liu et al., 2019). It has been demonstrated that an arbovirus can be replicated in multiple mosquito tissues including midguts (MGs), carcasses (CAs, tissues excluding MGs and SGs), and SGs, but the efficiency of replication in these tissues has not been fully evaluated (Bowers et al., 1995; King and Hillyer, 2012; Raquin and Lambrechts, 2017). Despite their small size, SGs are critical for arbovirus transmission, serving as the interface between the virus and the human host (Chowdhury et al., 2020; Liu et al., 2019; Ribeiro et al., 2016; Sanchez-Vargas et al., 2021). Although the exact number of viruses produced in one pair of SGs and released into saliva remains unclear, it is presumed to be substantial, as high viral titers are typically required to establish infection in the human host (Styer et al., 2007).

Mosquitoes possess a sophisticated immune system to manage arbovirus infections without succumbing to disease, maintaining viral loads that are sufficient for transmission to vertebrate hosts while simultaneously preventing harm to themselves (Cheng et al., 2016; Lee et al., 2019; Samuel et al., 2018; Sim et al., 2014). Perturbations of this antiviral defense, either through depletion or overactivation, have been shown to negatively impact arbovirus infection dynamics and transmission efficacy (Dong and Dimopoulos, 2023; Dong et al., 2022b; Jupatanakul et al., 2017; Merkling et al., 2023; Olmo et al., 2018; Samuel et al., 2023). The small interfering RNA (siRNA) pathway is the major antiviral defense against arbovirus infections in *Aedes* mosquitoes, where the RNase III enzyme Dicer 2 (Dcr2) recognizes and cleaves exogenous double-stranded viral RNA into 21-nucleotide siRNAs (Dong and Dimopoulos, 2023; Samuel et al., 2023). These exogenous viral siRNAs are then incorporated into the RNA-induced silencing complex (RISC) with the integration of dsRNA-binding protein called R2D2 and an endoribonuclease Argonaute 2 (Ago2). The guide strand of siRNAs directs the RISC to target the viral RNA complementary sequence, which will be cleaved by the Ago2, leading to the degradation of viral mRNAs. In addition to the core components, the siRNA pathway has multiple chaperon proteins such as TATA-binding protein-associated factor 11 (TAF11), Hop, Hsc70, Hsp83, and others, which contributes to RISC stability and enhance RNA splicing activities (Iwasaki et al., 2015; Liang et al., 2015).

While the siRNA pathway has been well characterized in the MGs and CAs, its role within the SGs remains largely unexplored (Bonning and Saleh, 2021; Franz et al., 2006; Liu et al., 2019; Olmo et al., 2018). The efficiency of this pathway in SGs and its impact on viral replication and transmission are unclear. Previous studies have demonstrated that tissue-specific overexpression of immune effectors such as *Dcr2, Loqs2, Hop* and *Dome* can significantly reduce viral replication in mosquitoes, although the extent of this effect varies between tissues and viral species (Dong et al., 2022b; Jupatanakul et al., 2017; Olmo et al., 2018). Despite these insights, the relative contributions of MGs, CAs, and SGs to overall vector competence remain poorly understood. Furthermore, it remains unknown whether simultaneously immune activation across multiple tissues could synergistically yield enhanced antiviral defenses.

In this study, we compared the expression of immune-related genes and titters of multiple arboviruses across different mosquito tissues, including MGs, CAs, and SGs. Our findings reveal that SGs have lower antiviral response and higher viral titers than CAs, suggesting reduced siRNA pathway activity in SGs. Overexpression of *Dcr2* in SGs, driven by a salivary gland-specific *Apyrase* promoter (Coates et al., 1999), significantly suppressed infections of both alphaviruses and orthoflaviviruses in SGs. Moreover, dual overexpression of *Dcr2* in both MGs and SGs resulted in a synergistic antiviral effect, effectively blocking the transmission of multiple arboviruses, including DENV, ZIKV, and MAYV. These results highlight the potential of targeting SG-specific immunity to disrupt arbovirus transmission and offer a promising genetic control strategy for reducing the global burden of arboviral infections.

## Results

### Mosquito midguts and salivary glands have lower expression of immune genes than carcasses

To investigate tissue-specific immune responses, we analyzed immune gene expression in midgut (MG), salivary glands (SGs), and carcass (CA) using publicly available transcriptomic datasets. Currently, there are three available RNA-seq datasets for *Ae. aegypti* SGs (Bonizzoni et al., 2012; Chowdhury et al., 2020; Ribeiro et al., 2016); however, only two were included in our analysis due to the availability of biological replicates, which are essential for robust statistical evaluation. Bonizzoni et al. generated RNA-seq data for the MGs, CAs and SGs, with two replicates per tissue (Bonizzoni et al., 2012). Chowdhury et al. sequenced SGs with two replicates (Chowdhury et al., 2020), while RNAseq data for CAs and MGs were retrieved from (Hyde et al., 2020). Transcriptomic analysis revealed significantly lower immune gene expression in MGs and SGs compared to CAs (Figure 1A-B, Table S1), suggesting a diminished immune response in these two tissues.

**Figure 1.**
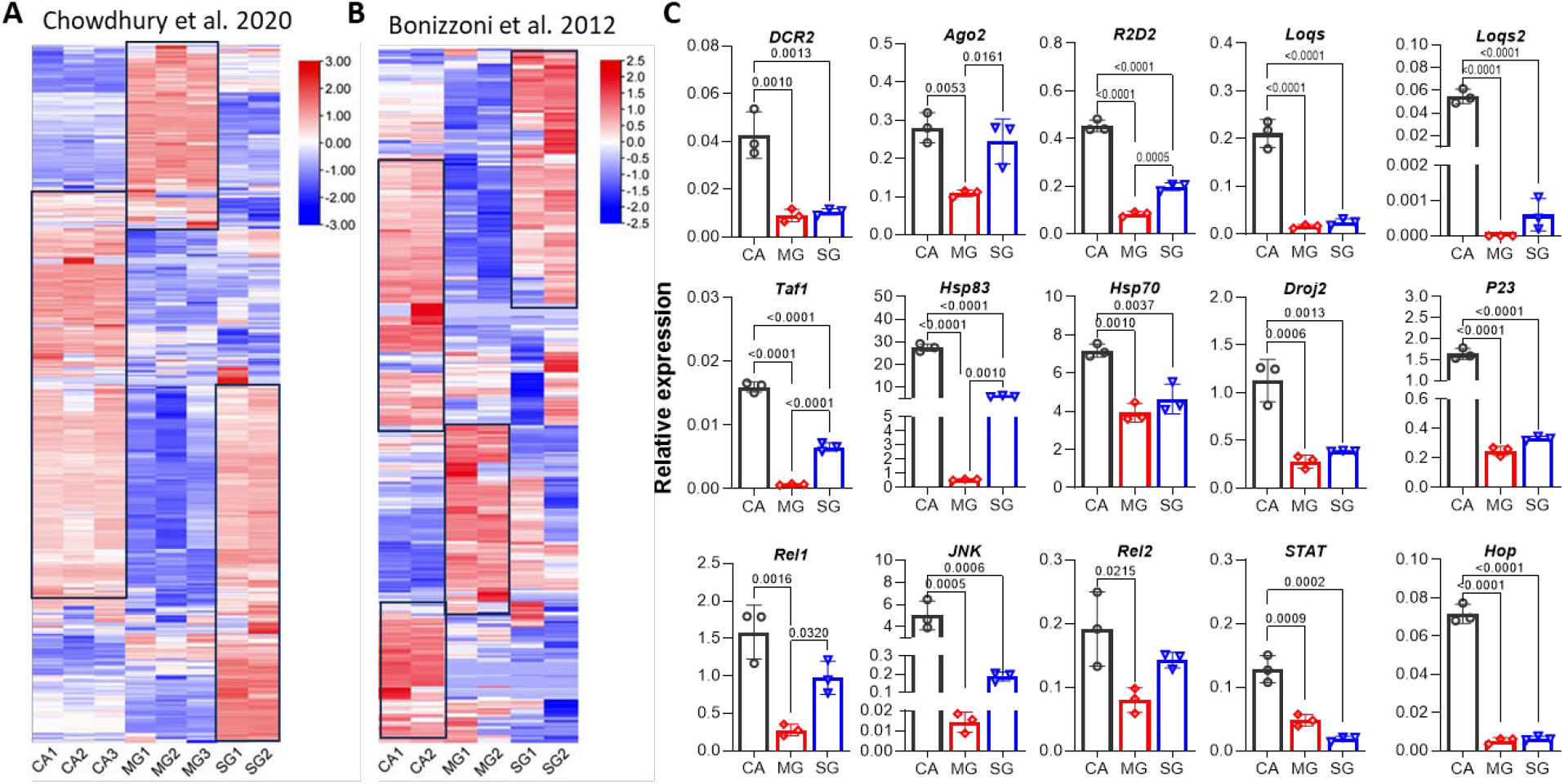
Tissue-specific expression of immune-related genes in different tissues of *Aedes aegypti*. **A-B**, Heatmaps showing transcriptome differences of immune genes among carcasses (CA), midguts (MG), and salivary glands (SG) by analyzing previously published RNA-seq datasets. Black boxes highlight regions of relatively high immune gene expression. Detailed gene expression data are provided in Supplementary Table 1. Color scale represents log2 fold change, with red indicating upregulation and blue indicating downregulation. **C**, qPCR analysis of core and chaperon genes of the siRNA pathway and classical immune pathway genes across tissues at 14 days post-ZIKV infection. Gene expression was normalized to *rpS7* and presented as relative expression to *rpS7*. Data are displayed as mean ± standard deviation (SD). *P* values were determined by one-way ANOVA. Dots indicate individual biological replicates (n=3), and bars indicate mean values.

To further investigate tissue-specific differences in antiviral defense, we assayed the expression of antiviral immunity genes in these tissues at 2 and 14 days post-ZIKV infection using qPCR. Core genes of the siRNA pathway were detected in all tissues at both time points; however, CAs exhibited significantly higher expression levels compared to MGs and SGs, particularly for *Loqs* and *Loqs2*, which were markedly reduced in MGs and SGs (Figure 1C and Figure S1). Similarly, chaperon genes (*Taf1, P23, Hsp83, Droj2* and *Hsp70*) and other antiviral immune genes (*Hop, JNK*, and *STAT*) exhibit significantly reduced expression in SGs and MGs compared to CAs (Figure 1C and Figure S1). These results, together with transcriptome analyses, support the hypothesis that SGs and MGs possess diminished antiviral defense compared to CAs, potentially contributing to their increased susceptibility to arbovirus infection.

### Salivary glands are highly susceptible to arbovirus infection

To investigate the impact of tissue-specific immune gene expression on viral infection, we analyzed viral load in CAs, MGs, and SGs following infection with two orthoflaviviruses (DENV2 and ZIKV) and one alphavirus (MAYV) using plaque assay. Given the difference in the extrinsic incubation period of orthoflaviviruses and alphaviruses in *Ae. aegypti*, tissues were dissected at 14 days post-infection (dpi) for DENV2 and ZIKV, and 7 dpi for MAYV, to ensure that the viruses had successfully infected and replicated in SG cells. The results showed that CAs exhibited significantly higher viral loads of DENV2 and MAYV compared to MGs and SGs, while MGs had a significantly lower virus load of ZIKV than CAs and SGs (Figure 2A).

**Figure 2.**
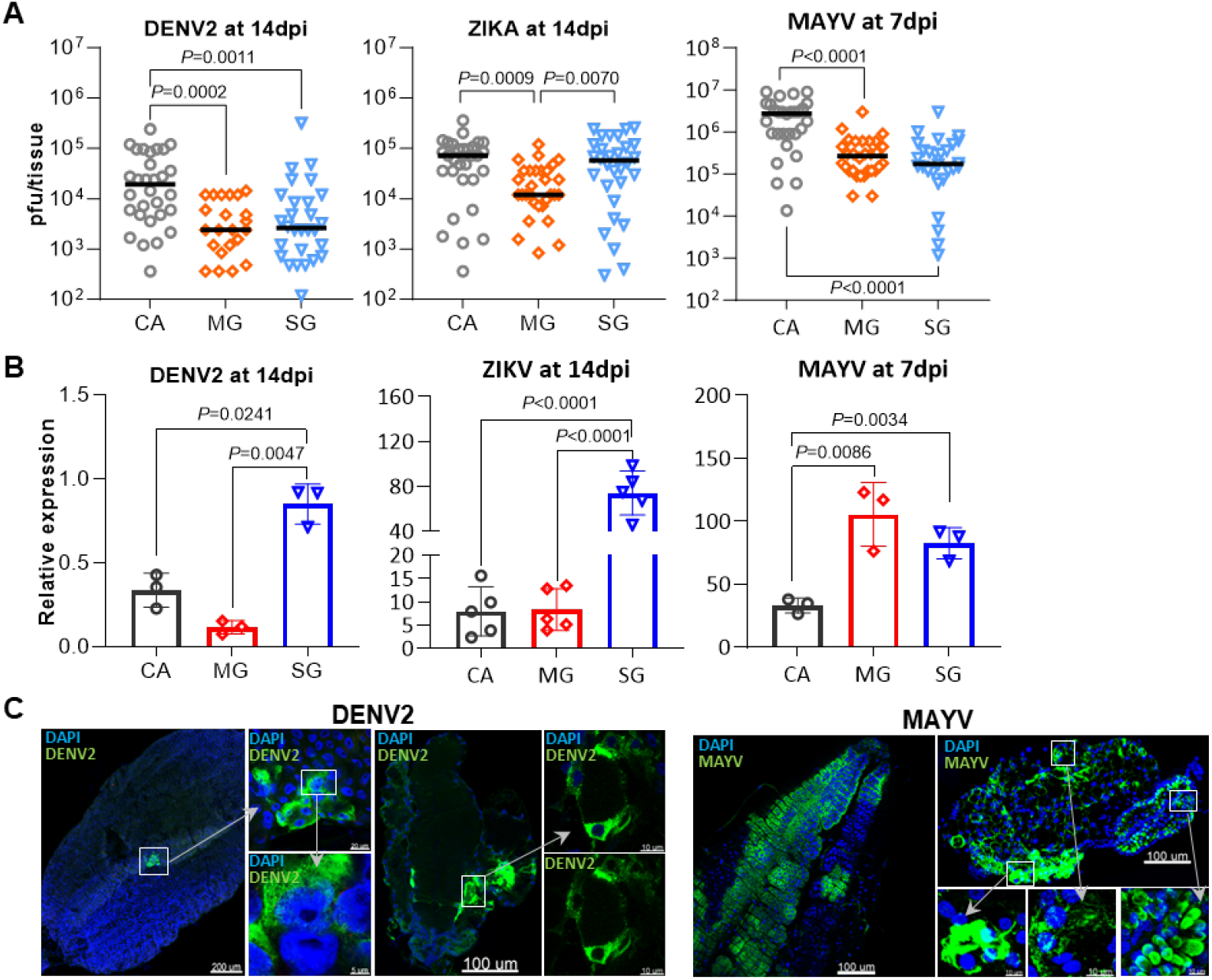
Comparison of viral titers in different tissues of *Aedes aegypti* infected by alphaviruses and orthoflaviviruses. **A**, Plaque assay measuring the titer of dengue virus 2 (DENV2) and zika virus (ZIKV) at 14 days post infection (dpi) and Mayaro virus (MAYV) at 7dpi in individual carcass (CA), midgut (MG) and a pair of salivary glands (SGs). Horizontal lines indicate medians. *P* values were determined by the Mann-Whitney test. **B**, qPCR detecting relative expression of DENV2 and ZIKV at 14dpi and MAYV at 7dpi in CAs, MGs and SGs. Viral RNA expression levels were normalized to the ribosomal S7 gene. Data are presented as mean ± SD. *P* values were determined by one-way ANOVA. **C**, IFA detecting infection patterns of DENV2 at 14dpi and MAYV at 7dpi in MGs and SGs. Specific monoclonal antibodies targeting DENV2 or MAYV (green) were used for virus detection, and nuclei were counterstained with DAPI (blue). The zoomed regions are highlighted with white boxes and arrows.

Considering the smaller size of SGs compared to CAs and MGs, we used qPCR to compare the relative virus titer in these tissues. The results showed that SGs exhibited higher relative virus titers (normalized to *rpS7* genes) of DENV2 and MAYV compared to CAs. SGs also had higher titers of two orthoflaviviruses compared to MGs, although this trend was not observed for the alphavirus (Figure 2B).

To further characterize infection patterns, we performed IFA on MGs and SGs. The analysis showed that the infection intensity of DENV2 in MGs was much lower than that in SGs. For MAYV, SGs also exhibited a higher intensity of infection than MGs, with the infection in MGs appearing more widespread, while SGs displayed infection restricted to specific regions (Figure 2C).

Collectively, these data indicate that SGs are highly susceptible to both orthoflaviviruses and alphaviruses, suggesting that they may be less refractory to arboviral infection compared to CAs.

### Overexpression of *Dcr2* in salivary glands effectively suppresses arbovirus infection

To investigate the impact of enhanced antiviral activity in SGs on arbovirus infection, we overexpressed *Dcr2* in SGs specifically using the *Apyrase* (*Apy*) promoter (Figure 3A). Four independent transgenic lines were established based on their fluorescent eye markers, and they showed different levels of resistance to DENV2 infection in SGs (Figure S2). The line (P1) with the strongest resistance to DENV2 infection was selected to establish a homozygous line (*ApyDcr2*). Expression of *Dcr2* was assayed in *ApyDcr2* females using qPCR, revealing a significant increase in *Dcr2* expression in SGs of *ApyDcr2* mosquitoes at 2-, 7-, and 14-days post blood-feeding (BF) compared to WT mosquitoes (Figure 3B).

**Figure 3.**
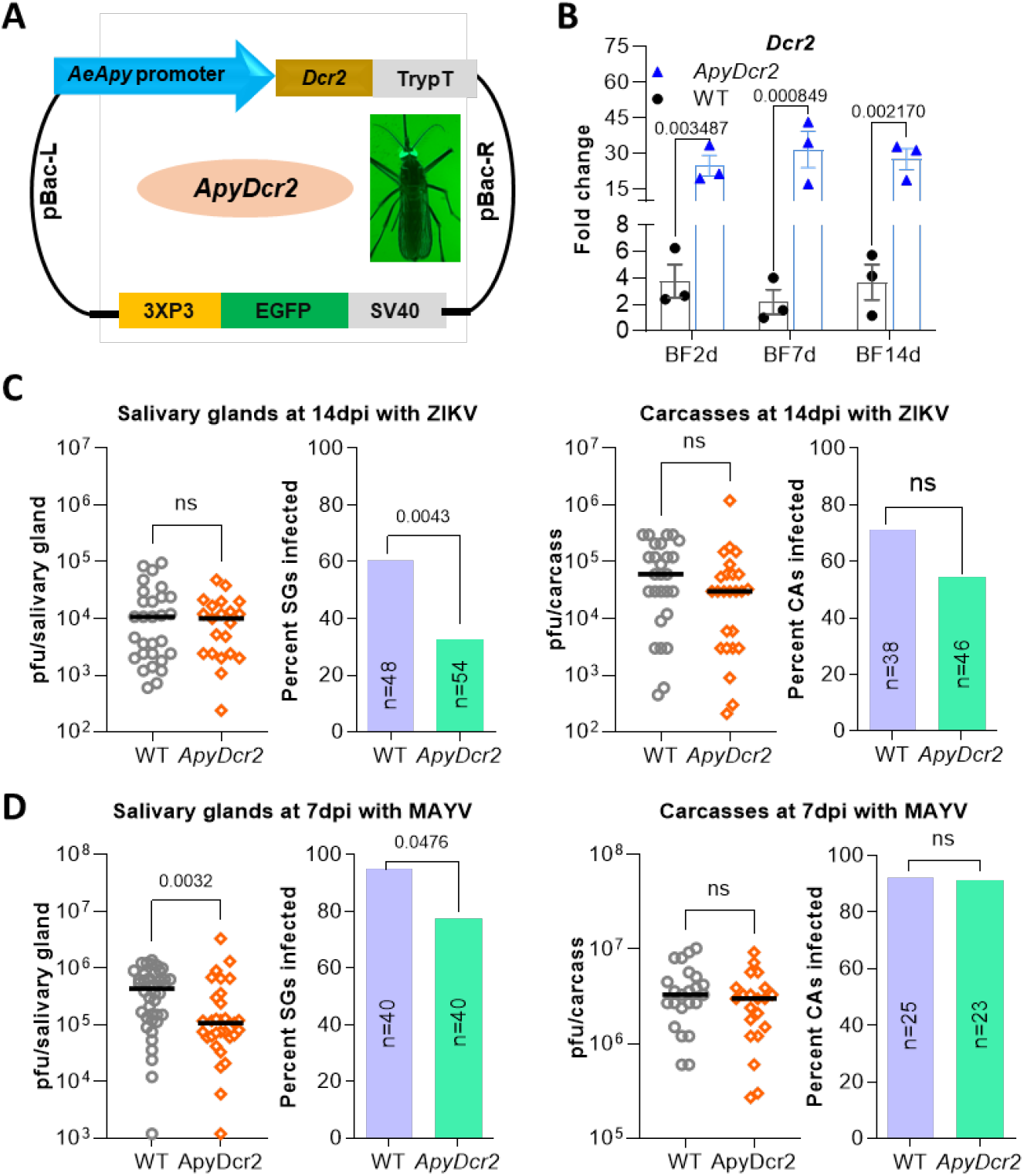
Generation of transgenic *Aedes aegypti* overexpressing *Dcr2* in salivary glands and examination of antiviral effect against ZIKV and MAYV infection. **A**, Scheme of the *ApyDcr2* construct for overexpressing *Dcr2* under the control of the salivary gland-specific *Apyrase* (*Apy*) promoter. Fluorescent green eyes in adult transgenic mosquitoes were used as a visual marker to identify positive individuals. Abbreviations: SV40, polyadenylation signal of simian virus 40 VP1 gene; 3xP3, eye tissue-specific promoter; TrypT: trypsin terminator sequence; pBac-L or -R, *piggybac* transposon elements left or right arms. **B**, Fold-change of transcript abundance of *Dcr2* in the salivary glands of the *ApyDcr2* transgenic line and wild-type line (WT) at different days post blood feeding (BF). Each bar represents a fold change of the *Dcr2* gene. The *rpS7* gene was used to normalize gene expressions, and error bars indicate the S.E. of the mean (shown as bars), with each dot indicating a value from each replicate, n=3. *P*-values for the transgenic line vs. the WT line were determined by using an unpaired t-test with GraphPad Prism. **C-D**, ZIKV (**C**) and MAYV (**D**) infection intensity (virus titer) and infection prevalence (numbers in the graphs indicate all tested mosquitoes) in the salivary glands and carcasses of females at 14 days post infection (dpi) of ZIKV and 7dpi of MAYV. Horizontal lines indicate medians. *P*-values for virus titer and infection prevalence were determined by a Mann-Whitney test and a Fisher’s exact test, respectively.

We subsequently compared viral titers and infection prevalence in SGs and CAs between *ApyDcr2* and WT mosquitoes at 14 dpi with ZIKV and 7 dpi with MAYV. No significant differences in viral titers or infection prevalence were observed in CAs between *ApyDcr2* and WT mosquitoes for either ZIKV or MAYV (Figure 3C-D). However, SGs from *ApyDcr2* mosquitoes displayed a significant reduction in infection prevalence for both ZIKV and MAYV, as well as a significant decrease in MAYV titter (Figure 3C-D).

In addition, there were no significant difference in key fitness parameters, including pupation time and rate, adult longevity (both females and males maintained on the sugar meal), body weight, blood-feeding percentage and efficiency, fecundity, and egg hatching rate, between *ApyDcr2* and WT mosquitoes (Figure S3). These results indicate that *Dcr2* overexpression in *SGs* does not impose measurable fitness costs on mosquito development, reproduction, or survival.

In summary, SG-specific *Dcr2* overexpression effectively suppresses both orthoflavivirus and alphavirus infection without compromising mosquito fitness.

### Overexpression of *Dcr2* in fat bodies has a mild effect on infection with DENV2 and no effect on MAYV

Given that transgenic overexpression of *Dcr2* in MGs (Dong et al., 2022b) or SGs significantly increases the mosquito’s resistance to arbovirus infection, we explored whether overexpression of *Dcr2* in CAs could confer similar refractoriness. To this end, we generated transgenic mosquitoes overexpressing *Dcr2* in fat bodies using *vitellogenin* (*Vg*) promoter (Jupatanakul et al., 2017), and three independent *VgDcr2* lines (P1, P2 and P3) were established (Figure 4A). qPCR analysis showed that P2 and P3 lines exhibited a significantly higher (2-to 3-fold) level of *Dcr2* expression compared to the WT at 24 hours post blood-feeding (Figure 4B), and a high level of *Dcr2* transgene was also observed in fat bodies of different *VgDcr2* lines but not in that of WT mosquitoes (Figure 4C). Further comparison of *Dcr2* transgene expressions in ovaries, fat bodies, and whole mosquitoes showed that the *Dcr2* transgene was predominantly expressed in the fat bodies, whereas endogenous *Dcr2* was expressed in all tissues (Figure 4D). However, the transgenic *Dcr2* expression levels in fat bodies were relatively low compared to endogenous *Dcr2* (Figure 4D), resulting in a smaller overall increase in *Dcr2* expression in *VgDcr2* mosquitoes compared to *ApyDcr2* and *CpDcr2* transgenic lines.

**Figure 4.**
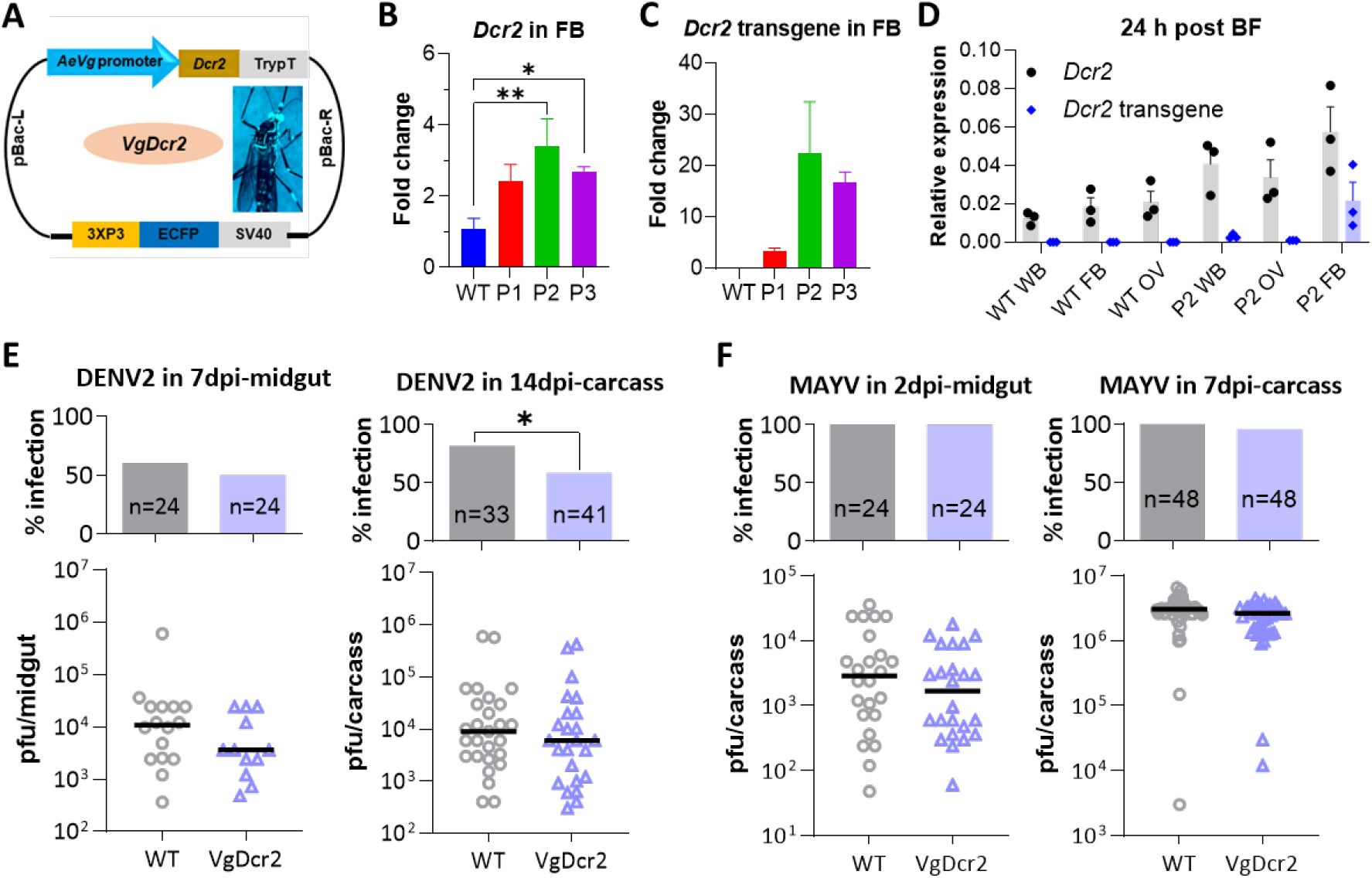
Generation of transgenic *Aedes aegypti* overexpressing *Dcr2* in fat bodies and examination of antiviral effect against DENV and MAYV infection. **A**, Scheme of *VgDcr2* construct overexpressing *Dcr2* under the control of the fat bodies-specific *vitellogenin* (*Vg*) promoter. Fluorescent blue eyes in adult transgenic mosquitoes were used as a visual marker to identify positive individuals. Abbreviations: sv40, polyadenylation signal of simian virus 40 VP1 gene; 3xP3, eye tissue-specific promoter; TrypT: trypsin terminator sequence; pBac-L or -R, *piggybac* transposon elements left or right arms. **B**, Transcript abundance of *Dcr2* in the fat bodies of different *VgDcr2* transgenic lines (P1, P2 and P3) and wild type (WT) at 24 hours post blood feeding (BF). Each bar represents the fold change of *Dcr2* gene relative to WT. Gene expression was normalized to the *rpS7* gene, and error bars represent the standard errors (S.E.) of the mean. **C**, Relative transcript abundance of *Dcr2* transgene in the fat bodies of *VgDcr2* transgenic lines and WT mosquitoes at 24 hours post blood-feeding (BF). Each bar represents a fold change of the *Dcr2* gene relative to the P1 line. **D**, Comparison of *Dcr2* and *Dcr2* transgene expression levels in the fat bodies, ovaries (OV) and whole bodies (WB) of the *VgDcr2* transgenic line (P2) and WT at 24 hours post BF. Each bar represents the relative expression of *Dcr2* or *Dcr2* transgene to rps17 gene. **E** and **F**, DENV2 and MAYV infection prevalence (upper panels) and infection intensity (virus titer, lower panels) in the midguts and carcasses at different days post infection (dpi). Horizontal lines indicate medians. Statistical significance was determined using an unpaired t-test for **B** and **C**, two-way ANOVA for **D**, and a Mann-Whitney test for virus titer or a Fisher’s exact test for infection prevalence for **E** and **F**. *P<0.05, **P<0.01.

*VgDcr2* (P2 line) and WT were subsequently infected with DENV2 and MAYV, and virus titer and infection prevalence were analyzed and compared in both MGs and CAs. For DENV2, there were no differences in viral titer or infection prevalence in midguts at 7 dpi, nor viral titer in carcasses at 14 dpi between *VgDcr2* and WT mosquitoes; however, a significantly reduced infection prevalence was observed in CAs of *VgDcr2* mosquitoes (Figure 4E). Conversely, no significant differences in viral titers or infection prevalence were detected for MAYV in either MGs (at 2 dpi) or CAs (at 7 dpi) between *VgDcr2* and WT mosquitoes (Figure 4F).

In conclusion, fat body-specific *Dcr2* overexpression resulted in a moderate reduction of DENV2 infection in CAs but had no effect on MAYV infection. These results suggest that *Dcr2* may not be a limiting factor for the siRNA pathway against alphavirus infection in CAs.

### Simultaneous overexpression of *Dcr2* in salivary glands and midguts efficiently blocks transmission of multiple arboviruses

To examine the effect of simultaneous *Dcr2* overexpression in SGs and MGs on arbovirus transmission, we generated a transgenic mosquito line (*CpApyDcr2*) by crossing *ApyDcr2* females (green eyes) with *CpDcr2* males (red eyes) (Figure 5). Following four generations of inbreeding, 90% of the offsprings had red and green, fluorescent eyes, and these mosquitoes were used for the arbovirus infection experiments. Two orthoflaviviruses, DENV2 and ZIKV, and two alphaviruses, MAYV and CHIKV, were tested in *ApyDcr2* and *CpApyDcr2* transgenic lines, with Liverpool strain serving as a WT control.

**Figure 5.**
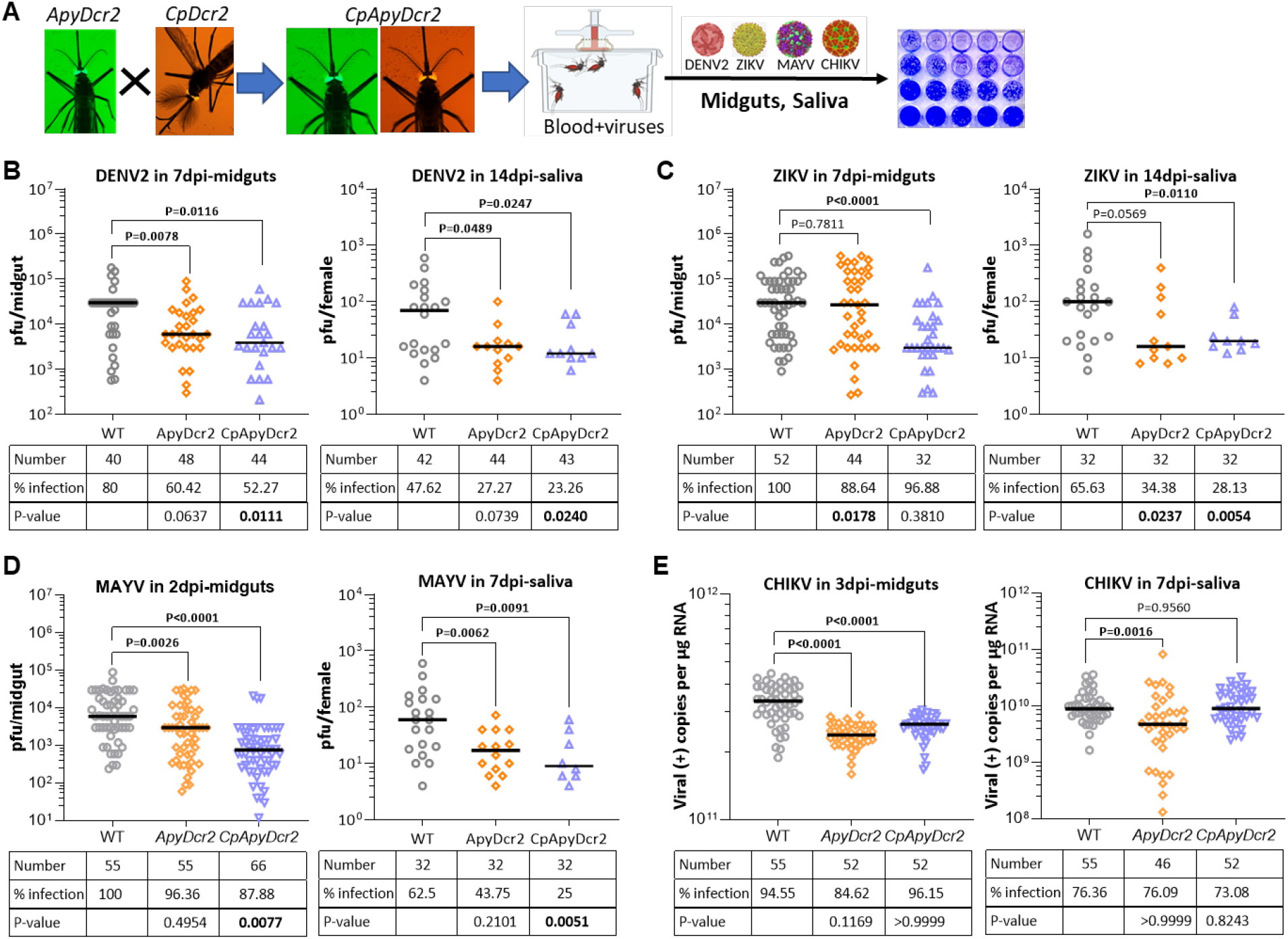
Effect of co-overexpression of *Dcr2* in *Aedes aegypti* salivary glands and midguts on arbovirus infection and transmission. **A**, Scheme of experiment design for generating transgenic mosquitoes that overexpress *Dcr2* in both midguts and salivary glands, followed by virus infection assay. *ApyDcr2* female mosquitoes (green eyes) were crossed with *CpDcr2* males (red eyes), and their offspring (*CpApyDcr2*) were screened for the presence of both green and red eyes. **B-E**, Virus titer and infection prevalence of DENV2 (**B**), ZIKV (**C**), MAYV (**D**) and CHIKV (**E**) in midguts and saliva of *ApyDcr2, CpApyDcr2* and *WT*. The tables below each graph summarize the total number of tested mosquitoes, percentage of infection, and *P* value for infection prevalence. Virus infection was determined by plaque assay for DENV2, ZIKV and MAYV and qPCR for CHIKV. Horizontal lines indicate medians. *P* values between transgenic and WT mosquitoes were determined by an unpaired two-sided Mann-Whitney test for virus titer or an unpaired two-sided Fisher’s exact test for infection prevalence.

The results showed that *CpApyDcr2* had significantly lower virus titers in MGs for all the tested arboviruses and in saliva for DENV2, ZIKV and MAYV (Figure 5B-E). Additionally, infection prevalence for DENV2, ZIKV, and MAYV was significantly reduced in both MGs and saliva of *CpApyDcr2* mosquitoes compared to WT controls (Figure 5B-E). While a marginal, non-significant reduction on infection prevalence for CHIKV was observed in saliva of *CpApyDcr2* (Figure 5), significantly lower CHIKV titers were observed in heads (including SGs) of *CpApyDcr2* mosquitoes (Figure S4). In addition, *ApyDcr2* mosquitoes displayed significant inhibition of virus titer for DENV2, MAYV and CHIKV in MGs and saliva, as well as reduced infection prevalence for ZIKV in MGs and saliva (Figure 5B-E).

These results demonstrate that dual overexpression of *Dcr2* in MGs and SGs significantly inhibits infection and replication of both orthoflaviviruses and alphaviruses, effectively blocking their transmission by mosquitoes. These findings suggest that concurrent expressions of the same antiviral effector in multiple tissues can synergistically enhance transmission-blocking efficiency within the same mosquito.

## Discussion

Arbovirus infection in *Ae. aegypti* progresses sequentially through the midgut, carcass, and salivary glands, with each tissue playing distinct roles in viral replication and/or transmission (Cheng et al., 2016; Franz et al., 2015; Sim et al., 2014). Our transcriptomic and qPCR analyses revealed a significantly lower expression of antiviral immune genes, including core components and chaperone genes of the siRNA pathway, as well as classical antimicrobial genes, in salivary glands and midguts compared to carcasses (Bonizzoni et al., 2012; Chowdhury et al., 2020; Hyde et al., 2020). Our findings align with previous studies that the lower expression of the siRNA genes and dsRNA-mediated silencing efficiency were observed in *Anopheles gambiae* salivary glands (Boisson et al., 2006). These results suggest that mosquito salivary glands exhibit a diminished antiviral immune response, potentially rendering them more permissive to viral infection and dissemination.

Infection assays further confirmed these observations, revealing that salivary gland cells exhibited higher viral titers compared to carcasses for both orthoflaviviruses (DENV2, ZIKV) and the alphavirus (MAYV), although carcasses displayed the highest overall viral loads. Additionally, immunofluorescence assays showed distinct infection patterns in different tissues, with salivary glands displaying focalized infection patterns, whereas midguts showed more diffuse viral distribution. Together with previous studies, these findings underscore the salivary glands’ critical role in arbovirus amplification and release (Bowers et al., 1995; Fiorillo et al., 2022; Raquin and Lambrechts, 2017; Salazar et al., 2007).

The reduced immune response in salivary glands and midguts may reflect an evolutionary trade-off, prioritizing physiological functions such as digestion and nutrient absorption in midguts and saliva production in salivary glands over robust antiviral response (Baron et al., 2010; Hixson et al., 2022; Sanchez-Vargas et al., 2021). In contrast, carcasses, comprising fat bodies, hemocytes, muscles, and other non-epithelial tissues, appear better equipped to mount a robust immune response (Cheng et al., 2016; King and Hillyer, 2012; Liu et al., 2019). Fat body tissues, analogous to the mammalian liver, are central to immune signaling and antimicrobial peptide production in mosquitoes, while hemocytes play key roles in phagocytosis and pathogen clearance. These immune components collectively contribute to systemic antiviral defense, reducing the viral burden in mosquitoes (Liu et al., 2019; Sim et al., 2014). Overall, these findings highlight the intricate balance mosquitoes maintain between immune defense and essential biological functions, shedding light on the tissue-specific roles in arbovirus replication, amplification, and transmission.

Importantly, the weaker antiviral defense observed in salivary glands suggests a promising target for genetic intervention to block arbovirus transmission. Indeed, overexpression of *Dcr2*, a core component of the RNAi pathway, in salivary glands resulted in a significant reduction in arbovirus transmission, emphasizing the potential of enhancing antiviral immunity in salivary glands to disrupt the arbovirus transmission cycle. In contrast, overexpressing *Dcr2* in fat bodies, the main tissues in carcasses, only moderately inhibited DENV2 infection and had no significant effect on MAYV infection. This discrepancy may be explained by differences in the baseline immunity and tissue-specific expression levels of transgenic *Dcr2*. Fat bodies express relatively high levels of endogenous *Dcr2*, and the additional transgenic expression might not result in a substantial increase in the overall siRNA-mediated antiviral activity above the existing threshold. Conversely, salivary glands, with inherently lower levels of endogenous *Dcr2*, showed a more pronounced antiviral effect when *Dcr2* expression was increased. Additionally, previous studies have demonstrated that *Dcr2* overexpression in midguts significantly increased antiviral defense (Dong et al., 2022b). These findings collectively highlight the importance of considering tissue-specific differences in immune dynamics when designing genetic interventions.

Targeting tissues with lower antiviral activity may represent a strategic approach to maximize the impact of engineered immunity and disrupt arbovirus transmission more effectively.

While we observed a significant increase in *Dcr2* expression in the salivary glands of *ApyDcr2* transgenic mosquitoes, we were unable to completely abolish viral infection. This may result from the *Apyrase* promoter being active only in a subset of salivary gland cells or from excessive *Dcr2* activity triggering anti-RNAi feedback mechanisms that may counteract the impact of siRNA pathway-mediated antiviral response (Blair, 2023; Moon et al., 2015; Pijlman et al., 2008; Poirier et al., 2018). The complex interactions between arboviruses and mosquito hosts also influence the role of *Dcr2* (Bonning and Saleh, 2021). Similar observations have been made in other transgenic mosquitoes engineered for resistance to *Plasmodium* or arbovirus infections (Dong et al., 2022a; Olmo et al., 2018; Wang et al., 2021; Williams et al., 2020). While these mosquitoes showed significantly increased expression of anti-pathogenic effectors, none achieved complete blockage of pathogen transmission, highlighting that genetic variation or epigenetic regulation may contribute to these outcomes, which has been observed in the natural mosquito population (Amarante et al., 2022; Lai and Wang, 2024; Lewis et al., 2023; Novelo et al., 2023). From a genetic control perspective, achieving robust and sustainable pathogen resistance in transgenic mosquitoes may require the identification and utilization of more tissue-specific promoters, a broader repertoire of antiviral effectors, and a strategic coordinated combination of multiple resistance mechanisms that are spatially and temporally targeted.

In addition, significant inhibition of DENV2, MAYV, and CHIKV was observed in the midguts of *ApyDcr2* transgenic mosquito lines. Previous studies have shown that the *Apyrase* gene is specifically expressed in mosquito salivary glands (Pala et al., 2024; Smartt et al., 1995), excluding the possibility of a direct effect of *Dcr2* overexpression in the midgut on viral inhibition. During blood feeding, mosquitoes ingest their own saliva (Klug et al., 2023; Luo et al., 2000; Pala et al., 2024), which contains proteins, miRNAs and viral siRNAs (Fiorillo et al., 2022). Notably, transgenic expression of *Dcr2* lacks a secretory signal peptide, preventing its secretion into saliva. This suggests that any viral inhibition observed in the midgut is unlikely to result directly from higher amount of *Dcr2* in the salivary glands. Therefore, further investigation is needed to elucidate the detailed mechanisms underlying these observations.

To evaluate the antiviral efficacy of engineered immunity in multiple tissues, we simultaneously overexpressed *Dcr2* in both midguts and salivary glands. This dual-tissue expression strategy achieved an even more pronounced antiviral effect, blocking transmission of multiple orthoflaviviruses (DENV2 and ZIKV) and alphaviruses (CHIKV and MAYV) more effectively than when *Dcr2* was overexpressed in salivary glands alone. However, antiviral resistance in *CpApyDcr2* mosquitoes seems to be dependent on arbovirus species, as evident by no difference in CHIKV infection density and prevalence in saliva. While we observed a significant reduction of CHIKV titer in heads, which includes salivary gland tissues. This discrepancy could be explained by differences in saliva collection/assay methods: filter paper collection for CHIKV genomic RNA preferred due to increased biosafety levels, vs forced salivation for DENV2, ZIKV and MAYV. In the former method, mosquitoes were allowed to salivate for 24 hours on filter paper, and viral load was measured using qPCR, capturing the cumulative virus load over 24 hours (Ledermann et al., 2024). Measurements of genomic RNA levels also tend to be higher than levels of infectious virus particles, as not all packaged virions will be infectious in a plaque assay conducted with mammalian cell lines. In contrast, the latter method involved forcing mosquitoes to salivate within one hour, followed by plaque assay to measure virus load, quantifying only infectious virions (Miller et al., 2021). As a result, the former method may show higher infection prevalence and viral titer compared to the latter method (Ledermann et al., 2024).

While our study provides significant implications for understanding mosquito-pathogen interactions and developing innovative strategies to combat mosquito-borne diseases, there are several considerations for future investigation. First, explore how these genetic modifications interact with other immune pathways, given the complexity of mosquito immunity (Lewis et al., 2023; Sim et al., 2014). Additionally, evaluating the efficacy of transgenic mosquitoes against field strains of arboviruses, rather than lab-adapted strains, would provide more applicable insights for real-world deployment. Finally, assessing the long-term stability and inheritance of the *Dcr2* transgene and its antiviral effects across multiple generations remains crucial for sustainable application.

In conclusion, our findings reveal the tissue-specific dynamics of antiviral immunity in mosquitoes and highlight the critical role of salivary glands in arbovirus transmission. By targeting the weak antiviral defenses in salivary glands through genetic modification, we can develop innovative and effective strategies to combat mosquito-borne diseases and mitigate their global impact.

## Materials and Methods

### Ethics statement

This study was conducted in compliance with the Guide for the Care and Use of Laboratory Animals of the National Institutes of Health and the recommendations of the institutional Ethics Committee (permit number M006H300). The protocol (permit #MO15H144) was approved by the Animal Care and Use Committee of Johns Hopkins University. Mice were used exclusively for mosquito rearing. Commercially sourced, anonymous human blood was utilized for mosquito virus infection assays; therefore, informed consent was not applicable.

All CHIKV infections were performed under biosafety level 3 (BSL-3) conditions at the Texas A&M Global Health Research Complex (TAMU, College Station, TX). The Institutional Biosafety Committee (IBC) approval number for this work is IBC2019-074.

### Mosquitoes

The *Ae. aegypti* Liverpool IB12 (LVP) strain and *CpDcr2* line (Dong et al., 2022b) were reared and maintained at 27°C under 85% relative humidity and a 12-hour light/dark cycle in the insectary of Johns Hopkins Malaria Research Institute. All transgenic mosquitoes were kept under identical conditions in a walk-in chamber. Mosquito larvae were fed with TetraMin tablets, while adult females were sustained on a 10% sucrose solution and blood-fed on ketamine-anesthetized Swiss Webster mice (Charles River Laboratories) for colony maintenance. For CHIKV infection, all mosquitoes were maintained at 28°C with 70–80% humidity and had continuous access to 10% sucrose solution.

### Bioinformatics analysis of RNA-Seq data

Two raw datasets for midguts, salivary glands and carcasses of uninfected mosquitoes were retrieved from NCBI. The accession number for the first dataset is PRJNA174376 (Bonizzoni et al., 2012), and the second dataset includes PRJNA609359 for midguts and carcasses (Hyde et al., 2020) and PRJNA533031 for salivary glands (Chowdhury et al., 2020).

Raw data were trimmed using Trimmomatic to remove adapters and low-quality reads, and clean reads were then aligned to the *Ae. aegypti* genome (VectorBase-68). Transcript abundance in each sample was quantified using FeatureCounts with gene annotations from VectorBase. Differentially expressed (DE) genes between midguts, salivary glands and carcasses were identified using DESeq2 at Usegalaxy (https://usegalaxy.org/). Gene ontology (GO) and fold enrichment of the DE genes were performed using ShinyGO (http://bioinformatics.sdstate.edu/go/), and heatmaps were generated using SRplot (https://www.bioinformatics.com.cn/srplot).

### RT-qPCR

Midguts (MG), salivary glands (SG), and carcasses (CA, defined as the whole body excluding MG and SG) were dissected from mosquitoes in sterile PBS on ice. Each sample consisted of 10 MG, 30 SG, or 10 CA. The samples were homogenized in 300 μL TRIzol (Invitrogen, Carlsbad, CA), and total RNA was extracted following the standard protocol. First-strand cDNA was synthesized using the QuantiTect reverse transcription kit (Qiagen, Hilden, Germany). qPCR amplifications were performed with gene-specific or virus-specific primers (Table S2) and ABI SYBR Green Supermix on the StepOnePlus real-time PCR system (Applied Biosystems, Warrington, UK), as previously described (Dong and Dimopoulos, 2023). The relative abundance of the gene transcripts or virus load was normalized to the ribosomal protein S7 gene and calculated using the 2−^ΔΔ^CT or 2−^Δ^CT comparative method. Each sample had at least three independent biological replicates.

### Immunofluorescence assays (IFA)

Midguts and salivary glands were dissected from WT mosquitoes at 14 dpi for DENV2 and at 7 dpi for MAYV, respectively. Samples were fixed overnight at 4°C in 100 μl of 4% paraformaldehyde (Sigma) in a PCR tube, and then were permeabilized with 0.2% Triton X-100 in PBS containing 1% bovine serum albumin (TBB). After three washes with PBST (PBS with 0.1% Triton X-100), samples were incubated overnight at 4°C with anti-MAYV E2 antibody (Sigma) or mouse hyperimmune ascitic fluid specific for DENV2 (Xi et al., 2008), followed by incubation with the secondary antibody, AlexaFluor 488 goat anti-mouse IgG (Invitrogen), in TBB for 2 h at 37°C in the dark. DAPI (4′,6-diamidino-2-phenylindole) was added to stain the nuclei. Samples were then visualized and photographed using a Zeiss LSM700 confocal microscope at the Johns Hopkins University School of Medicine Microscope Facility.

### Generation of transgenic *Ae. aegypti* mosquitoes using *piggyBac* transposase

The *Ae. aegypti Apyrase* and *vitellogenin* promoters were cloned from LVP mosquitoes using specific primers and inserted into the pENT plasmid. The *Dcr2* gene was amplified from the pMos-CpDcr2 plasmid (Dong et al., 2022b). Subsequently, the *Dcr2* and the *Apyrase* (or *vitellogenin*) promoters were assembled into the *piggyBac*-eGFP (or *piggyBac*-eCFP) backbone plasmids using gateway cloning (Jupatanakul et al., 2017). The final plasmid was sequenced through, and the sequence is available upon request. The plasmid was then chemically transformed into *E. coli* DH5α cells and purified using the EndoFree Maxi Prep kit (Qiagen, Germantown, USA). The *piggyBac* transposase mRNA was synthesized via *in vitro* transcription (IVT) using a HiScribe® T7 ARCA mRNA Kit (New England Biolabs, NEB) and purified with RNAClean XP beads (Beckman). The IVT template was prepared by PCR amplification of a plasmid containing the *piggyBac* transposase with specific primers (Hacker et al., 2023).

Germ-line transformation was carried out following our previously published method (Dong et al., 2022b). Briefly, *piggy*Bac transposase mRNA and donor plasmid were mixed with nuclease-free water to a final concentration of 300 ng/μL of each. This mixture was injected into freshly laid LVP eggs using a FemtoJect Express (Eppendorf). The injected eggs were hatched in distilled water at 3 days post-injection, and the resulting larvae (G0) were reared according to standard mosquito-rearing protocols. The surviving adults were mated to the opposite-sex LVP mosquitoes, and their offspring (G1) were screened for fluorescent eye markers. The positive G1 pupae were sexed, and separate pools were established. The heterozygous mosquitoes were backcrossed with LVP mosquitoes for four generations, followed by inbreeding for another four generations to establish homozygous lines for downstream experiments.

### Assessment of fitness parameters

Fitness parameters, including larval development, adult body weight, female blood-feeding capacity, fecundity, egg hatchability, and longevity, were compared between transgenic and WT mosquitoes following our previously published methods (Dong and Dimopoulos, 2023; Dong et al., 2022b). Briefly, the larval development was assayed in a container with 50 larvae, and the pupation time was recorded daily. The body weight was determined with five females using a fine balance. The blood-feeding capacity was determined with 10 females in cup, and blood-meal intake per mosquito was determined by measuring the difference in group weight before and after blood-feeding.

The fecundity and egg fertility were assayed using the agarose-based method (Tsujimoto and Adelman, 2021). The blood-fed females laid eggs on 2% agarose in a 24-well plate, and the eggs were hatched in the same plate. Eggs and hatched larvae in each well were photographed and counted. Male and female longevity was monitored in 16-oz paper cups containing 20 mosquitoes each. Mortality was recorded daily for up to 60 days.

### DENV2, ZIKV and MAYV infection and transmission assay

DENV serotype 2 (DENV2; New Guinea C strain), ZIKV (Cambodia, FSS13025) and MAYV (Iquitos, IQT, gifted from Dr. Alexander W.E. Franz at University of Missouri) were propagated in C6/36 cells as described previously (Dong and Dimopoulos, 2023; Dong et al., 2016). Virus-infected cell culture medium was harvested at 5 days post-infection (dpi) for DENV2 and ZIKV, and 3 dpi for MAYV. The media were mixed with an equal volume of commercial human blood supplemented with 10% human serum and 10 mM ATP to prepare the infectious blood meal. Approximately 60 one-week-old transgenic or wild-type (WT) females were starved overnight in a 32-oz paper cup. The mosquitoes were then fed for 30 minutes on a glass artificial feeder containing the blood-virus mixture. Fully engorged females were sorted in a 4°C cold room, transferred to a fresh cup, and maintained on a 10% sterile sucrose solution in the insectary. The viral infection was performed at ACL3 in the insectary at Johns Hopkins Malaria Research Institute.

At 7 dpi with DENV2 and ZIKV, and 3 dpi with MAYV, midguts (MGs) were dissected from each individual female in sterile PBS on ice. Similarly, salivary glands (SGs) and carcasses (CAs) were dissected at 14 dpi with DENV2 and ZIKV, and 7 dpi with MAYV. Samples were transferred to Eppendorf tubes containing 300 µl DMEM with glass beads and stored at −80°C. Samples were homogenized using a Bullet Blender (Next Advance, Inc., Averill Park, NY) for 3 minutes at speed 3, followed by centrifugation at 7,000 × g for 3 minutes at 4°C. Filtered supernatants (50 µl for MGs, 100 µl for SGs, and 20 µl for CAs) were used for plaque assays on BHK-21 cells (DENV2) or Vero cells (ZIKV, MAYV) in 24-well plates. The plates were incubated for 3 days (MAYV), 4 days (ZIKV), or 5 days (DENV2), followed by staining with 1% crystal violet to visualize plaques. Plaques were counted under a microscope to determine viral titters. Each experiment included at least two biological replicates, with more than 15 individual females per replicate.

Viral transmission was assessed using forced salivation at 7 dpi for MAYV and 14 dpi for DENV2, following previously established protocols (Dong et al., 2021). Mosquitoes were knocked down in a cold room, and wings and legs were removed on ice. The proboscis of each mosquito was inserted into a 10 μl pipette tip containing 2.5 μl of Immersion Oil Type B (Sigma). Mosquitoes were allowed to salivate for 30 minutes, and saliva droplets were observed under a microscope. Collected saliva, along with the oil, was transferred into PCR tubes containing 50 μl DMEM (10% FBS, 1% penicillin/streptomycin) on ice. Samples were immediately frozen on dry ice and stored at −80°C for viral titration using plaque assays.

### CHIKV infection and transmission assay

CHIKV (strain 99659) stocks were propagated in Vero E6 cells (ATCC C1008) cultured in Dulbecco’s Modified Eagle Medium (DMEM; Gibco, Thermo Fisher Scientific, Waltham, MA) supplemented with 10% heat-inactivated fetal bovine serum (FBS) and 1% penicillin/streptomycin. Virus concentrations were quantified using plaque assays on Vero cells. Three-to five-day-old females were starved for 24 hours before feeding. The virus was diluted in defibrinated sheep’s blood (Colorado Serum, Denver, CO) to a final concentration of 1 × 10^6^ PFU/mL. Blood meals were offered for one hour using a Hemotek feeding system (Discovery Workshops, Accrington, UK) fitted with an artificial membrane. Fully engorged mosquitoes were separated into two groups: (1) groups of 20 mosquitoes were maintained per container for midgut infection assays, and (2) individual mosquitoes were placed into 12-well plates with mesh overlays for transmission assays (Ledermann et al., 2024).

Midguts were dissected at 3 dpi, transferred into 2 mL tubes containing 600 µL RAV1 buffer (MACHEREY-NAGEL, Allentown, PA), and homogenized for two sequential cycles using a TissueLyser II bead mill (Qiagen, Germantown, MD) at 25 Hz for 30 seconds per cycle. For transmission analysis, mosquito heads and corresponding sucrose-feeding filter papers were placed in separate 2 mL screw-cap tubes with 400 µL RAV1 buffer each and homogenized under the same conditions.

Homogenized samples were immediately frozen at −80°C until further processing. RNA was extracted using the MACHEREY-NAGEL NucleoSpin RNA Virus kit according to the manufacturer’s instructions. cDNA was synthesized using Superscript IV reverse transcriptase (Thermo Fisher Scientific, USA) with RNA inputs of 300 ng (MGs), 100 ng (heads), and 30 ng (filter papers). Strand-specific quantitative real-time PCR (ssq-PCR) was performed using strand-specific primers and probes as previously described (Plaskon et al., 2009). Samples with a Ct value >35 were considered CHIKV-positive. Viral copy numbers were quantified using a standard curve derived from the CHIKV NSP1 sequence.

### Statistical analysis

All statistical analysis was performed using GraphPad Prism 10. Differences in the virus titer, fecundity, and hatching rate between transgenic and WT mosquitoes were assessed by an unpaired two-sided Mann–Whitney test, while infection prevalence by a two-sided Fisher’s exact test. *P* values for differences in larval development and survival rate between transgenic and WT mosquitoes were determined using the Gehan-Breslow-Wilcoxon test and the log rank (Mantel-Cox) test, respectively. The significance of differences in gene expression was analyzed using one-way ANOVA, and differences in body weight and blood-feeding capability were assessed using Student’s *t*-test.

## Acknowledgments

This work was supported by National Institutes of Health / National Institute of Allergy and Infectious Disease R01AI141532 and the Bloomberg Philanthropies. The founders had no role in study design, data collection and analysis, decision to publish, or manuscript preparation. We thank Johns Hopkins Malaria Research Institute Parasitology Core facilities for providing the naive human blood for arbovirus infection assays. We also thank Ms. Aliyah Silver, a rotation Ph.D student, for her assistance with *ApyDcr2* construct generation, and Dr. Deborah McClellan for editorial assistance. The illustrations in the figures were created with BioRender.com.

## Author contributions

S.D., Y.D., and G.D. conceived and designed the experiments. S.D., J.C., and Y.Z. performed the experiments. S.D., J.C., Y.D., K.M., and G.D. analyzed the data, and S.D., J.C., and G.D. wrote the paper, with input from all authors. All authors commented on the manuscript.

## Data and materials availability

All data needed to evaluate the conclusions in the paper are present in the paper and/or the Supplementary Materials. Complete sequence maps and plasmids and *Ae. aegypti* transgenic lines are available upon request to G.D.

## Supplementary figures and table

**Figure S1** qPCR analysis of core and chaperon genes of the siRNA pathway and classical immune pathway genes across tissues at 2 days post-ZIKV infection. Gene expression was normalized to *rpS7* and presented as relative expression to *rpS7*. Data are displayed as mean ± standard deviation (SD). *P* values were determined by one-way ANOVA. Dots indicate individual biological replicates (n=3), and bars indicate mean values.

**Figure S2 Infection intensity (virus titer) and infection prevalence (numbers on the graph indicate all tested mosquitoes) in salivary glands of different pools of *ApyDcr2* transgenic lines at 14 days post infection (dpi) with DENV2**. Horizontal lines indicate medians. Statistical significance was determined by a Mann-Whitney test for infection intensity or a Fisher’s exact test for infection prevalence. *P<0.05, **P<0.01. WT, wild-type strain (Liverpool); P1, P2, P3 and P8: different pools of transgenic lines overexpressing *Dcr2* in salivary glands.

**Figure S3 Fitness parameters of *ApyDcr2* mosquitoes as compared to those of WT mosquitoes. A**, larval development and pupation rate. **B**, longevity of females and males. **C**, body weight of newly emerged adult females and males. **D**, blood-feeding capability of females. **H**, fecundity (eggs laid per mosquito) and fertility. The *P* value was determined by a Gehan-Breslow-Wilcoxon test for (**A**) and a log-rank (Mantel-Cox) test for (**B**). Statistical significance between *ApyDcr2* and WT was determined by an unpaired two-sided *t*-test for (**C-D**) and an unpaired two-sided Mann-Whitney test for (**E**). ns, not significant.

**Figure S4 Virus titer and infection prevalence in heads of WT, *ApyDcr2* and *CpApyDcr2* females at 7 days post infection (dpi) with CHIKV**. The tables below the graph summarize number of tested mosquitoes, percentage of infection and *P* value. Virus titter was determined by qPCR. Horizontal lines indicate medians. *P* values between transgenic and WT mosquitoes were determined by an unpaired two-sided Mann-Whitney test for virus titer or an unpaired two-sided Fisher’s exact test for infection prevalence.

**Table S1 Expression of immune genes in carcasses (CAs), midguts (MGs) and salivary glands (SGs) by analyzing previously published RNA-seq datasets**.

**Table S2 Primer sequences**.

