## Supplementary Figures for "Enhancing Tissue-Specific Antiviral Immunity to Disrupt Arbovirus Transmission by Mosquitoes"

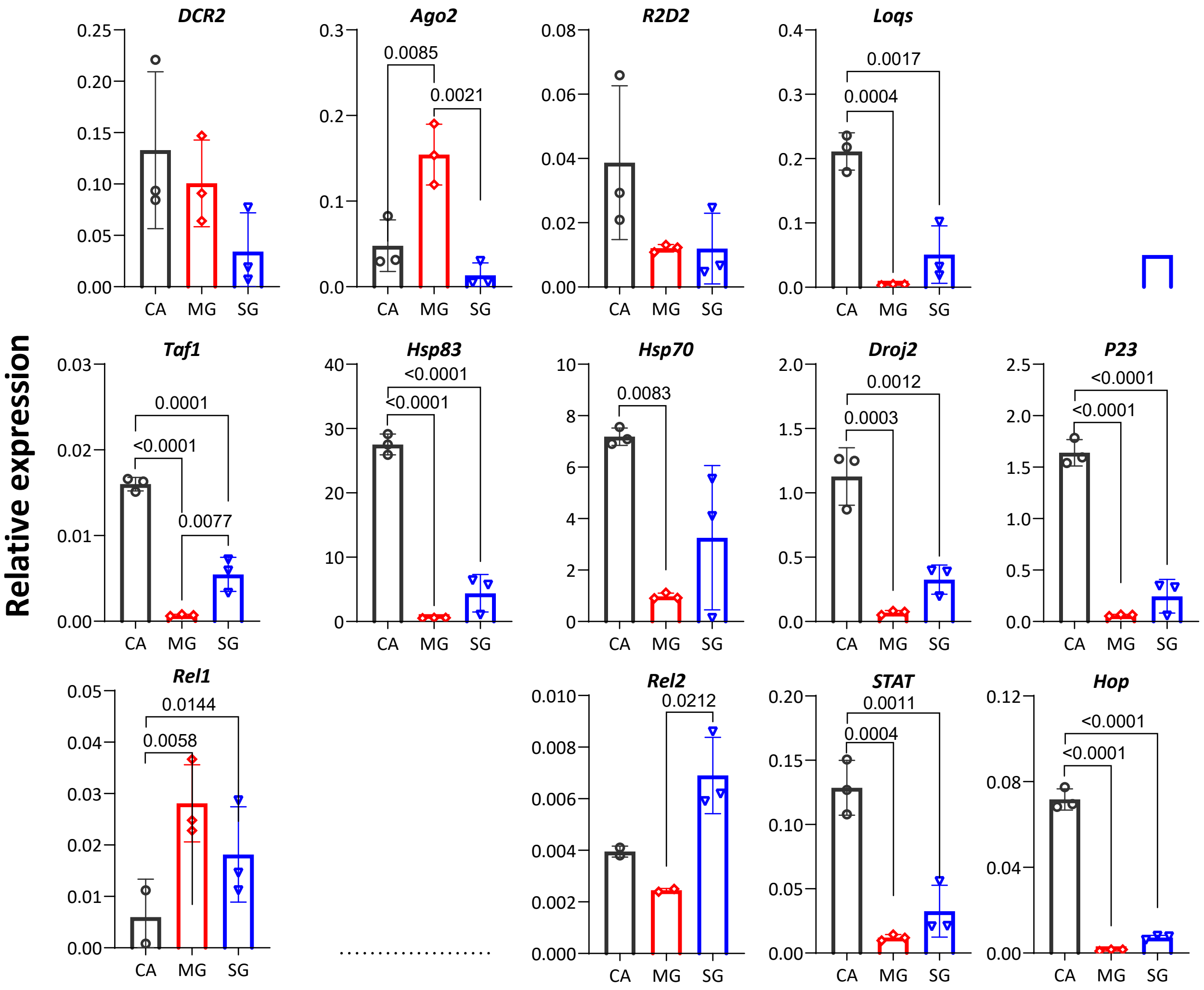
Relative expression
DCR2
Ago2
0.0085
0.0021
R2D2
Loqs
0.0017
0.0004
Taf1
0.0001
<0.0001
0.0077
Hsp83
<0.0001
<0.0001
Hsp70
0.0083
Droj2
0.0012
0.0003
P23
<0.0001
<0.0001
Rel1
0.0144
0.0058
Rel2
0.0212
STAT
0.0011
0.0004
Hop
<0.0001
<0.0001
CA MG SG

**Figure S1**

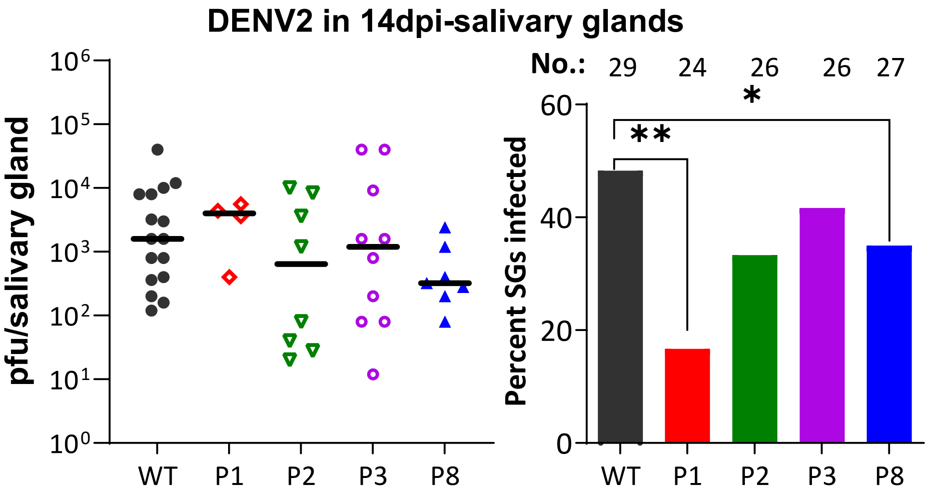
DENV2 in 14dpi-salivary glands
pfu/salivary gland
10^6
10^5
10^4
10^3
10^2
10^1
10^0
WT
P1
P2
P3
P8
No.: 29 24 26 26 27
Percent SGs infected
60
40
20
0
**
*
WT
P1
P2
P3
P8

**Figure S2**

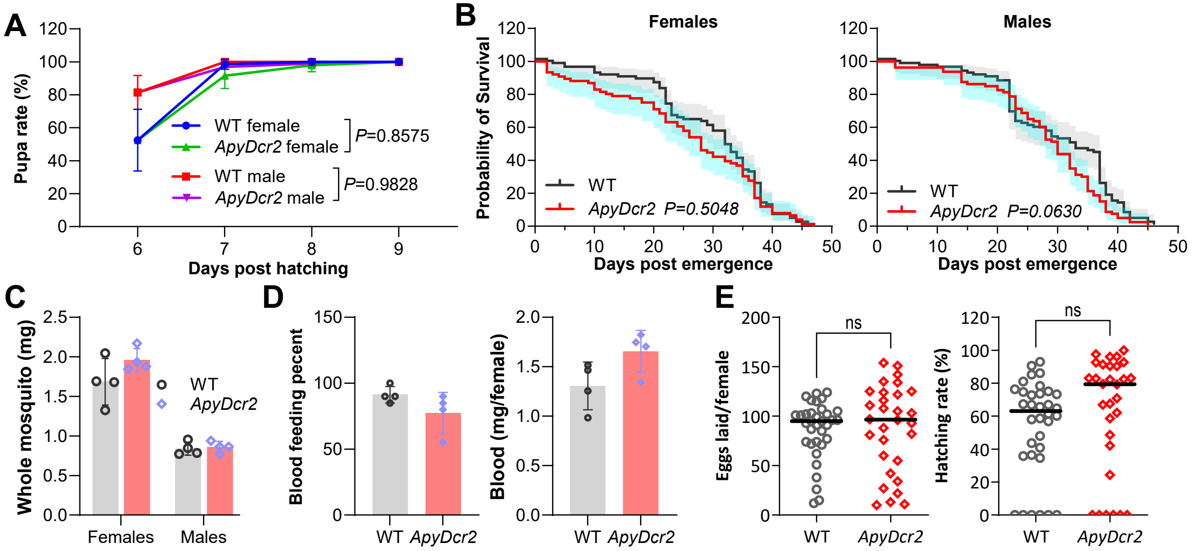

A
Pupa rate (%)
Days post hatching
WT female
ApyDcr2 female
P=0.8575
WT male
ApyDcr2 male
P=0.9828
B
Females
Males
Probability of Survival
Days post emergence
WT
ApyDcr2 P=0.5048
ApyDcr2 P=0.0630
C
Whole mosquito (mg)
Females
Males
WT
ApyDcr2
D
Blood feeding pecent
Blood (mg/female)
E
Eggs laid/female
Hatching rate (%)
ns


Figure S3

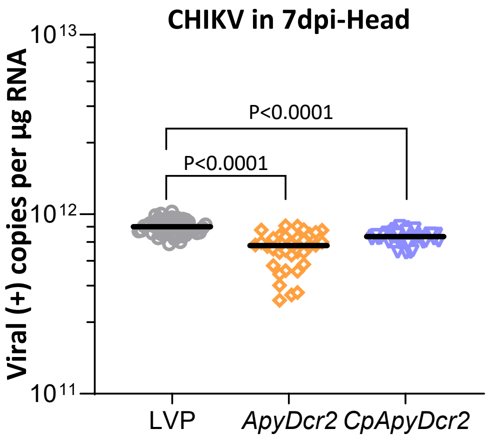


| | | | |
|---|---|---|---|
| Number | 55 | 37 | 52 |
| % infection | 90.91 | 91.89 | 96.15 |
| P-value | | >0.9999 | 0.4386 |

**Figure S4**

### Table S2 Primers

| Genes/Viruses | Primer Name | Sequence (5' to 3') | Purpose |
|---|---|---|---|
| AeApyrase (AAEL006347) | AeApyF_XhoI_PBS | CCGCTCGAGAGAATTCAGTCAACTCTGT | Transgenic construct cloning |
| | AeApyF_AvrII_PBS | CCGCCTAGGGTTTTCTTAACTAACAATTTTGCC | |
| | AeApyF_seq | AGTGCTTTGGTGCTTATTTGGT | |
| AeVgA (AAEL010434) | AeVgF_XhoI_PBS | CCGCTCGAGCACCGAATTCCACCACCAGG | |
| | AeVgF_AvrII_PBS | CCGCCTAGGGTCTTCAAGTATCCGGCAGC | |
| AeAgo2 (AAEL017251) | Ago2_F | CGAGATGATTAGAGATCTGC | qPCR |
| | Ago2_R | ATGGCACGAAGTTCTATGG | |
| AeDcr2 (AAEL006794) | Dcr2_R | CAACGCTTTCAGTCAAACGA | |
| | Dcr2_F | ATTGATCCCCCAAAAAGACC | |
| AeR2D2 (AAEL011753) | R2D2_F | GACCTACCGGGAACTCATCA | |
| | R2D2_R | GATCAGGGTGCATTTGTCCT | |
| AeLoqs (AAEL008687) | AaeLoqs_RT_F | AGAAATCGGTGAGCAAGCAG | |
| | AaeLoqs_RT_R | GCGGAAAGTTGGCTCGTGGAC | |
| AeLoqs2 (AAEL013721) | AaeLoqs2_RT_F | CGGTGTTGCTTTCTGGGATA | |
| | AaeLoqs2_RT_R | CATGGCTTCGGGACAGTAAA | |
| AeDroj2 (AAEL005165) | AaeDroj2_880F | GGCCAGGATCTGATCATGCA | |
| | AaeDroj2_1006R | GCTTCAAGACTTCACCCGGA | |
| AeHsc70 (AAEL019403) | AaeHsc70_1375F | CTCCACTGTCGCTCGGTATC | |
| | AaeHsc70_1493R | GGGTTGGTTGTCCGAGTAGG | |
| Hsp83 (AAEL011708) | AaeHsp83_1577F | TCCGCTTCTGGAGACGAGTA | |
| | AaeHsp83_1695R | AAGGCCGAGTTCTTGACCTG | |
| Taf11 (AAEL023506) | AaeTaf11_189F | TTCCACTCGTTGCTCTCGTC | |
| | AaeTaf11_271R | TGCAGCTGAGTTGGTTCGAT | |
| AeHop (AAEL012553) | AeHop_F | CCGGACTTTATCGAGCTGTC | |
| | AeHop_R | ATCTGGTTCACTCCGTCGTC | |
| AeRel2 (AAEL007624) | AeRel2_F | GCTCAGTGCTACCGTGGGAAAC | |
| | AeRel2_R | CCCCAGTCCTGCCAGATCTC | |
| AeRel1 (AAEL007696) | AeRel1_F | TGGTGGTGGTGTCCTGCGTAAC | |
| | AeRel1_R | GGATGCCCAGGTTGCTGAAGG | |
| AeStat (AAEL020559) | AeStat_F | CAACCTGCAGTATCCGGTTT | |
| | AeStat_R | GAACCACTCCCAGAAGGTGA | |
| AeJNK (AAEL008634) | AeJNK_F | GGTGATACCGAGTTTGAGGTTC | |
| | AeJNK_R | CGACATTCTGCTGGGTGATT | |
| Dcr2 transgene | AeVg5-UTR-F1 | CACACAATCGGAACAGCTGC | |
| | AeDcr2-R1 | CTCCGGTCGGCAAGTAGATG | |
| AeP23 (AAEL014943) | AaeP23_419F | ATCAACCGGTGCATCGAGTT | |
| | AaeP23_550R | CGAGCCCTCATCCTTCCATC | |
| MAYV | MAYV_F | CAAATGTCCACCAGGCGAAG | |
| | MAYV_R | GTGGTCGCACAGTGAATCTTTC | |
| ZIKV | ZIKV_F | AGCAACATGGCGGAGGTAAG | |
| | ZIKV_R | CTGTCCACTAACGTTCTTTTGCAGA | |
| DENV2 | DENV2_F | TCCCTTCCAAATCGCAGCAACAATG | |
| | DENV2_R | CGTTCTGTGCCTGGAATGATG | |
| AeS7 (AAEL009496) | rpS7_F | GCAGACCACCATTGAACACA | |
| | rpS7_R | CACGTCCGGTCAGCTTCTTG | |
| CHIKV | CHIKV R T tag T | GGCAGTATCGTGAATTCGATGC GTCCGCCCTTTGTCTACATGA | Strand-specific for CHIKV |
| | CHIK F T | aataaatcataaGACGCAGAAACGCCCACATT | |
| | Tag T | aataaatcataaGGCAGTATCGTGAATTCGATGC | |
| | CHIKV probe | TGCTTGCACACTGACGT | |
